# Phylogeographic Inference Using Approximate Likelihoods

**DOI:** 10.1101/025353

**Authors:** Brian C. O’Meara, Nathan D. Jackson, Ariadna Morales, Bryan C. Carstens

## Abstract

The demographic history of most species is complex, with multiple evolutionary processes combining to shape the observed patterns of genetic diversity. To infer this history, the discipline of phylogeography has (to date) used models that simplify the historical demography of the focal organism, for example by assuming or ignoring ongoing gene flow between populations or by requiring *a priori* specification of divergence history. Since no single model incorporates every possible evolutionary process, researchers rely on intuition to choose the models that they use to analyze their data. Here, we develop an approach to circumvent this reliance on intuition. PHRAPL allows users to calculate the probability of a large number of demographic histories given their data, enabling them to identify the optimal model and produce accurate parameter estimates for a given system. Using PHRAPL, we reanalyze data from 19 recent phylogeographic investigations. Results indicate that the optimal models for most datasets parameterize both gene flow and population divergence, and suggest that species tree methods (which do not consider gene flow) are overly simplistic for most phylogeographic systems. These results highlight the importance of phylogeographic model selection, and reinforce the role of phylogeography as a bridge between population genetics and phylogenetics.

---

Phylogeographic investigations operate at the scale where population-level processes begin to form phylogenetic patterns. As such, the field can act as a bridge between population genetics and phylogenetics (Avise et al. 1987), and provide valuable information about the early stages of speciation (Moritz 1994). While early phylogeographic studies were conducted largely from a phylogenetic perspective (Crandall and Templeton 1996; Sullivan et al. 2000), the incorporation of coalescent theory (Kingman 1982) has provided a theoretical basis for incorporating population level processes such as gene flow, genetic drift, and population size change into empirical investigations. This expansion of the evolutionary processes considered by phylogeography has improved the discipline immensely, with recent studies providing clear examples of the importance of processes such as range expansion and gene flow (e.g., Khatchikian et al. 2015; Weir et al. 2015). However, increasing the complexity of the historical models considered by phylogeographic research has an associated cost; in general, analytical solutions are not available for these models, and thus complex computational machinery such as Markov chain Monte Carlo (MCMC) are required to estimate parameters that quantify processes such as gene flow.

Researchers can choose among a wide range of software packages that implement powerful phylogeographic models; however, due to their inherent complexity, most of these methods impose limits on the parameter space under consideration. For example, the classic island model (Wright 1931) can be used to estimate gene flow using several methods. The popular program Migrate-*n* (Beerli and Felsenstein 2001) allows researchers to estimate rates of gene flow among *n* populations, but assumes that allele coalescence among populations is due to migration (i.e., it does not consider the temporal divergence among populations). This assumption is counter to those of species tree methods which estimate the branching history among populations, but under the assumption of no gene flow (see Edwards 2009). Because many phylogeographic investigations are concerned with both processes, so researchers have turned to isolation-with-migration (IM) models (Nielsen and Wakeley 2001), particularly IMa2 (Hey and Nielsen 2007; Hey 2010), which considers multiple populations. Such models approach the phylogenetic models that have long been applied to phylogeographic datasets, and may be particularly well suited to incipient phylogenetic systems (Hey and Nielsen 2004). However, IMa2 still imposes an important limit on the historical demography of the focal system; it does not infer the relationships among lineages. Given theoretical findings that demonstrate how gene flow can decrease the accuracy of species tree inference (Eckert and Carstens 2008; Leaché et al. 2014), an ideal method would estimate gene flow in addition to the pattern and timing of population isolation.

Phylogeographic methods derive signal from patterns of genetic variation inherent to the empirical data. Parameters estimated from the data are thus contingent on the parameterization of the model used to estimate a particular set of parameters, and consequently conflicting inferences can follow from analysis of the same data using different models (i.e., differing sets of parameters). Consider the example of *Myotis lucifugus*, a vespertillionid bat distributed throughout most of North America. Several subspecies of this bat have been described, and species delimitation analyses indicate that these subspecies are independent evolutionary lineages (Carstens and Dewey 2010; Fig. S1). However, analysis of these data using an *n*-island model (Migrate-*n*) support gene flow among 3 of the 4 subspecies (Table S1), while analyses using an IM model produce substantial estimates of gene flow among diverging lineages (Carstens and Dewey 2010). Thus, *M. lucifugus* subspecies are inferred to be independent evolutionary lineages if species tree models are used to analyze the data, or populations within a species that exchange alleles when an *n*-island model is used to analyze the data. Parameter estimates using an IM model suggest that both processes (i.e., gene flow and population divergence) are important, but this also suggests the possibility that estimates of lineage phylogeny could be mislead by not accounting for gene flow. While assessing the extent of gene flow among populations has long been a critical aspect of speciation research (Dobzhansky 1937; Mayr 1963), results such as these are difficult to resolve unless the statistical fit of the underlying models can be evaluated. Absent a framework for evaluating model fit, researchers have been forced to reconcile conflicting parameter estimates on a *post hoc* basis (Koopman and Carstens 2010).

In this report, we introduce a novel method (PHRAPL) that calculates the approximate likelihood of a large number of demographic models given the data, and demonstrate that it provides a suitable framework for assessing the statistical fit of the models commonly used in phylogeographic research. PHRAPL compares the topology of gene trees estimated from empirical data to those simulated under various demographic models. It then approximates the probability of the data given those demographic models by calculating the proportion of times that simulated gene tree topologies match the empirical topologies (O’Meara 2010), and adopts a multiple model inference framework (Burnham and Anderson, 2004) to quantify the support for each model in the comparison set. When applied to empirical systems such as *Myotis*, it enables researchers to identify *which* parameters are essential to estimate. PHRAPL is also likely to be useful as a tool for data exploration, particularly in systems that lack previous phylogeographic investigation.

To date, phylogeographic inference has been almost entirely based on a qualitative interpretation of parameters that were estimated using models that were selected based on the intuition of the researchers. Consequently, it is not clear if and how incorrect phylogeographic model choice has misled researchers. To explore this question, we reanalyze recently published data (Table S2) and compare the demographic models selected by PHRAPL to those chosen by researchers.

## Methods

### Description of the phylogeographic approximate likelihood method.

PHRAPL inputs gene tree topologies (without branch lengths) estimated from empirical data, as well as an association file that maps sampled individuals to populations. It assumes that recombination within genes can be ignored and that gene trees from different genes are effectively unlinked. The maximum number of free parameters (which determines the size of the set of possible models) must also be defined, either by the user or automatically. PHRAPL then creates a list of the possible demographic histories that can describe these populations. All models contain some combination of parameters that describe the time of population coalescence (*t*) and/or the migration rate (*M* = 4*Nm)* among some number of populations. Models with any combination of migration rates are possible; for example, one possible model could parameterize gene flow from population A to B as parameter *M*_1_, gene flow from population C to B and C to A as *M*_2_, and set all other migration rates set to zero. In addition, users can apply filters to simplify model space: for example, one can limit models to a maximum of two different population sizes, consider only models in which all populations coalesce, and so forth. While any restriction of model space is an exchange of generality for computational efficiency, such restrictions allow for flexible incorporation of existing knowledge, and enables researchers to use PHRAPL in either a completely agnostic manner or as a tool to evaluate specific biogeographic hypotheses. The current implementation of PHRAPL assumes constant population size during a given time interval (i.e., between population splitting events), but future implementations could relax this assumption to allow exponential population growth or decline. Once the complete list of models is specified by PHRAPL, each is converted into a command for the program ms (Hudson 2002), which is then used to simulate gene trees under the various models with particular numeric values of parameters. Log-likelihoods are approximated from the proportion of simulated topologies that match the observed ones, described in more detail below.

Rather than a full optimization search of parameter space, we present in this paper analyses that were conducted using a grid of parameter values. This was found in initial explorations to be more efficient than optimization (Fig. S2) and provides a better sense of the confidence intervals of parameter values than can be achieved with typical numerical optimization. Note that both a grid and continuous parameter optimization are available as options to users, although we recommend seeding optimization searches with values obtained from a preliminary grid search. Preliminary searches using a coarse grid (i.e., with large increments between proposed values) may be used to construct finer grids for subsequent analyses. In addition, if the optimal value is found to be at the extreme edge of a grid, further searches should be conducted to include more extreme values. Although the grid is composed of discrete values, parameter estimates from a given model are obtained by model averaging across the grid, and thus parameter estimates are continuous and not confined to taking on values included in the grid. Because the size of the grid rapidly expands with additional parameters, the coarseness of the grid must be balanced with desired model complexity, and it will be computationally difficult to search over using either a grid or optimization for models with large numbers of parameters (e.g. > 10).

PHRAPL is written in R, and is available at https://github.com/bomeara/phrapl.

### Strategies for increasing the efficiency of calculating approximate likelihoods of demographic models

One obvious challenge for PHRAPL is effectively searching the large set of possible gene trees. For just seven sampled alleles, there are 10,395 binary gene trees; this number increases to >10^21^ trees when 20 alleles are sampled and would exceed the square of the number of atoms in the universe when more than 100 alleles are sampled. On the surface, this would seem to preclude a method that calculates probability based on the proportion of gene trees that match a given history, but PHRAPL uses several strategies to circumvent this difficulty.

First, when comparing the empirical gene tree to the simulated gene trees, PHRAPL assumes that samples from within a population are interchangeable, and then corrects for this assumption. This allows PHRAPL to switch labels within a population: if the observed gene tree is (((A1,A2),A3),(B1,C1)), where letters indicate populations, and numbers indicate alleles sampled from that population, the simulated gene tree (((A3,A2),A1),(B1,C1)) matches the observed gene tree except for the switching of the labels within population A. Thus, when scoring matches, PHRAPL automatically considers all possible assignments of individual labels within populations and corrects for the number of such samplings (for example, in this case, there are six possible assignment permutations of labels A1, A2, and A3, so these are all tried and a match is counted as 1/6 of a possible match, as the expectation is that 1/6 of the simulations would result in a perfect match). This removes the stochastic factor of intrapopulation labeling on detecting matches, and results in a more efficient matching algorithm, particularly as the number of samples increases.

Second, although there may be millions of possible gene trees under a given demographic history, these do not occur at equal probability. For example, in cases of small populations that have been isolated for many generations, congruence between the gene tree and population tree occurs at a relatively high frequency (Hudson and Turelli 2003), so many gene trees will have the same interpopulation branches, leaving only intrapopulation disagreements. This non-uniform distribution of gene tree probabilities (Degnan and Salter 2005) enables a far more efficient PHRAPL inference: the main differences between models will come in the relative probabilities of fairly common trees, rather than in the relative probabilities of trees in the tail, so PHRAPL can achieve reasonable degrees of accuracy in assessing model fit with a feasible number of simulations.

Third, we implement subsampling of individuals within putative populations, which has been shown to be an effective strategy for estimating species trees from phylogeographic data (Hird et al. 2010). Even with full resolution, having tens to hundreds of samples within a population may not provide much more information than having fewer samples because most coalescent events occur very recently in the tip populations, but will require far greater computation time. PHRAPL randomly samples individuals from all populations (with number of samples per population and number of replicates specified by the user) and then analyzes these subsampled gene trees against the model set. Multiple sampling strategies are possible, although preliminary explorations have shown that subsampling three or four alleles per population yields the best balance between adequate information content and computational efficiency.

Finally, given that the approximate likelihood is calculated by counting matches, with a finite number of simulated trees there is some probability that none of them have matched a particular observed tree. This produces a point estimate of the likelihood of the model equal to zero, or a log likelihood of negative infinity. In practice, we know that all gene topologies have a nonzero, albeit sometimes very small, probability, but having one or more negative infinities in the log likelihood prevents the calculation of the overall AIC. Put another way, our estimate of the probability of a gene tree naturally comes from the maximum likelihood estimate of the probability of successes in a binomial distribution, but this estimate prohibits calculation of the approximate likelihood if it is exactly zero. We thus incorporate this small bit of prior information. Absent data, we expect the probability that an observed tree will match a given random tree is just one over the number of possible random trees with the same number of tips. We chose to give the prior a small weight: though we simulate thousands of trees, by default (this can be adjusted by the user) the prior only has as much weight as 100 additional simulated trees (*nEq*). In practice, this gives the approximate likelihood a tiny nudge away from exactly zero in the case of no matches, making the log likelihood finite, while having little effect in cases where there are observed matches. Our overall estimate of the likelihood of a gene tree *G* given the model (including parameters) *M* is not simply the number of matches (*nMatch*) divided by the number of simulated trees (*nSim*) but rather comes from a beta-binomial distribution:

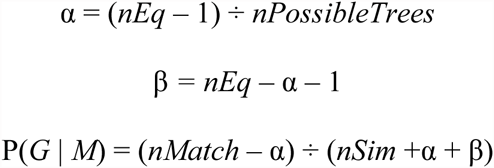

The effect of this change is relatively minor. Given 20 taxon gene trees, 10,000 simulated trees, and five gene trees that match, respectively, 1, 2, 0, 3, 0 times, this changes the likelihood estimate for each gene tree from 0.000100, 0.000200, 0.000000, 0.000300, and 0.000000, to 0.000099, 0.000198, 1.195e^-24^, 0.000297, and 1.195e^-24^, resulting in finite log-likelihoods and calculable approximate likelihoods.

### Demographic histories and model space.

One challenge to phylogeographic model selection is defining the complexity of the model space. The possible number of topologies for *n* sampled populations is slightly higher than the number of resolved and unresolved phylogenetic trees for the same number of OTUs (the difference comes from allowing histories with at least some non-coalescing populations as in a typical island population model). This number grows factorially with *n*, and can be large, even with only a few free parameters. For example, an empirical dataset with 3 populations, up to 2 population divergence events, and 1 migration rate would have 1,462 possible histories. To reduce the complexity of model space, constraints can be added; for example users can specify that population merging is to occur only between geographically proximate populations. At most, there will be *n*-1 possible free parameters for divergence times, 2*n*-2 possible population sizes at nodes (including terminal nodes) (not 2*n*-1, as the initial population size is fixed in all the models), 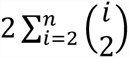 possible migration rates, and 2*n*-1 possible population growth rates. This explicit focus on the number of free parameters has an auxiliary benefit: since the number of possible parameters scales with the number of loci and the number of samples, users should explore tradeoffs in resource allocation and may be discouraged from proposing overly-complex demographic scenarios, a common problem in phylogeography (Knowles and Maddison 2002).

While PHRAPL is not essentially a Bayesian approach, it shares some similarities with approximate Bayesian computation (ABC; Beaumont et al. 2002), which has also been applied to phylogeographic model selection (e.g., Fagundes et al. 2007; Peter et al. 2010). In the absence of an analytical solution, both rely on data simulated under specific demographic scenarios, and evaluate these data by comparing them to the observed data. However, PHRAPL uses likelihood rather than a Bayesian criterion. It thus seeks the model (i.e,. demographic history + parameter estimates) that maximizes the probability of the data, rather than integrating over parameter space to calculate posterior probability. In ABC, prior beliefs about the distribution are required for parameter values (in some cases, these could be set to uninformative priors), and in some cases the priors can affect the final results (e.g., Oaks et al. 2013). PHRAPL does not use priors, but does require some specification of the region of parameter space to explore. PHRAPL also differs from ABC in how it evaluates similarity between the simulated and the observed data. ABC approaches typically use a variety of summary statistics and require specification of an epsilon value to describe how close a simulated value has to be to an empirical value to count as a “match”. PHRAPL uses discrete gene topologies in effectively the same way, as a summary statistic, but does not require specification of an epsilon value because topology is a discrete parameter. We have observed that log-likelihoods calculated using PHRAPL are generally good approximations of analytical log-likelihood estimates (Fig. S3), at least for those demographic models which have analytical solutions. Finally, PHRAPL is explicitly tailored for model selection. It calculates AIC scores for each model (Akaike 1973) and then applies information theory (Burnham and Anderson 1998) to assess the relative support of these models given a particular dataset. This enables model selection from a large set of demographic models, an objective that is empirically difficult when using ABC for model choice (Pelletier and Carstens 2014).

### Analyses with simulated data

We tested the performance of PHRAPL using simulated datasets for which the underlying history is known. We focused on four demographic histories (Fig. 1) each consisting of three populations: an isolation only (IO) model, isolation with migration (IM) model, *n*-island migration only (MO) model, and a mixed (MX) model which was intermediate to IM and MO models and included one population coalescent event, with migration between some, but not all populations. For IO and IM models, we considered three divergence depths (shallow, medium, and deep), yielding eight histories (Fig. 1). Under each history we simulated gene trees using ms for five different dataset sizes (1, 5, 10, 50, and 100 loci), with 20 replicates for each treatment (for a total of 480 datasets). Parameter values span an order of magnitude of migration rates and divergence times and were chosen to reflect the breadth of variation commonly observed within phylogeographic datasets.

**Fig. 1.**
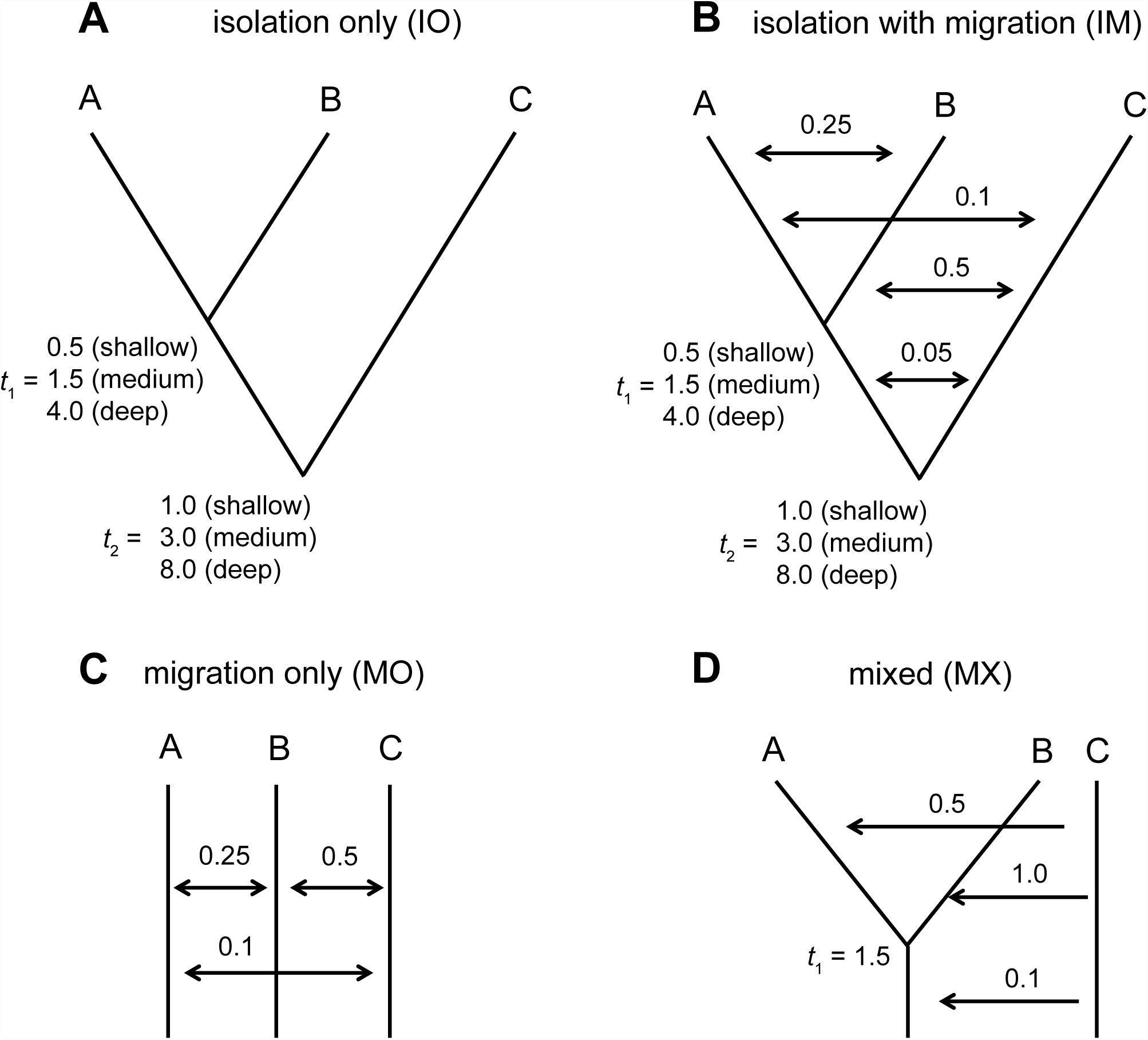
Histories used for simulation testing. Four types of histories were simulated, each involving three populations: A: isolation only models, which exhibit two collapse events, *t*_1_ and *t*_2_ (three time depths were simulated; times are given in units of 4*N*); B: isolation with migration models (migration rates above arrows are given in units of 4*Nm*); C: a migration only model; and D: a mixed model, which includes one collapse event and migration in some, but not all directions.

We analyzed these simulated datasets against two different model sets. First, we assembled a small model set (11 models) containing (i) the full true underlying models (four models), (ii) a set of simplified IM, MO, and MX models (three models, which are identical to the full true models except that they include only a single shared migration rate), and (iii) IO and IM models that reflect alternative branching histories (four models). Secondly, we analyzed simulated datasets against a large model set consisting of a filtered set of all possible demographic models that include one, two, or three free parameters (maxK = 3). To reduce model space we assumed a shared, constant population size across populations, set the maximum number of migration parameters to one, and considered only models in which migration parameters were symmetric between populations (e.g., if there is migration from population A to population B, there must also be migration from B to A). For three populations, the three available free parameters could thus be allotted to zero, one, or two possible population coalescence events and/or to one possible migration rate, in all possible combinations. After adding in three missing full models from the small model set, this yielded a total of 81 models. We also repeated all the above analyses (for 10 replicates and for only three of the five dataset sizes) using genealogies inferred from sequence data, which we evolved along all simulated genealogies using Seq-Gen (Rambaut and Grassley 1997; Figs. S4-S5). These analyses were designed to assess how the stochastic process of mutation influences PHRAPL inferences.

For each dataset, model, and value combination in the specified parameter grid, we simulated 10,000 gene trees. For the grid, we considered seven values of time to population coalescence (*t*) (0.30, 0.58, 1.11, 2.12, 4.07, 7.81, and 15.00) and six values of migration rate (*M*) (0.10, 0.22, 0.46, 1.00, 2.15, 4.64). For each simulation, the approximate log-likelihood of a dataset under a given model was calculated using the mean log-likelihood across subsampled trees for a given locus, and summing these across loci. We performed PHRAPL analyses on 10 iterative subsamples of each dataset, subsampling 4 individuals per population (resulting in 12 tip trees).

### Analyses with empirical data

Data were kindly provided by the corresponding authors of 19 recently-published phylogeographic investigations (Barker et al. 2012, Camargo et al. 2012, Harrington and Near 2012, Hung et al. 2012, Jackson and Austin 2012, Muir et al. 2012, Werneck et al. 2012, Carstens et al. 2013, Dhami et al. 2013, Duvernell et al. 2013, Fernandez-Mazuecos and Vargas 2013, Hamback et al. 2013, Leaché et al. 2013, Reilly et al. 2013, Satler et al. 2013, Talavera et al. 2013, Tsai and Carstens 2013; Wielstra et al. 2013, Giaria et al. 2014, Halas and Simons 2014; see Table S2 for specifications of included datasets). We selected studies (i) that were published or in press between January 2012 and October 2013; (ii) that used *BEAST (Heled and Drummond 2010) or IMa2 (Hey 2010), two of the most commonly applied methods for inferring evolutionary parameters of interest in phylogeographic datasets; (iii) that used sequence data from multiple loci that were either available to us from the authors or from public databases; (iv) that analyzed their data using three defined lineages (to allow for the same, moderately sized model set to be used for all datasets); and (v) sampled four or more individuals per population. If a dataset lacked an outgroup sequence, we obtained one from GenBank. If needed, datasets were aligned using MUSCLE (Edgar 2004). For each dataset and locus, we estimated a maximum likelihood tree using the rapid hill-climbing algorithm (10 replicate searches) and GTRGAMMA model implemented in RaxML-HPC v7.2.6 (Stamatakis 2006).

We analyzed each empirical dataset using the larger (81) model set used for the simulated data, except we removed the three models that contained more than one migration rate (for a total of 78 models). 200 subsample iterations were used for each dataset, with three individuals per population subsampled for each iteration (i.e., nine tip trees), as this yielded better AIC consistency across replicate runs of these datasets than was observed by subsampling four (Figs. S6-S7). The number of trees simulated per cycle was set to 100,000. The parameter grid was identical to that used for the simulated datasets, except that we added one additional migration rate (4*Nm* = 10). Parameter estimates for mtDNA were scaled to account for their ¼ effective population size.

In addition, we analyzed the *Myotis* dataset (Carstens and Dewey 2010) that contained four populations (one corresponding to each subspecies) using 216 total models. These models were filtered to include only those with a fully resolved population history, but varied in topology and presence and direction of migration.

## Results and Discussion

### Performance with simulated datasets

At moderate to deep levels of divergence, PHRAPL is generally accurate at identifying the ’true’ model (i.e., the model used to generate the data), although in some cases, this is contingent upon sampling many loci (Fig. 2). In cases where the generating model does not have the highest AIC score, this model is often ranked second-best and is surpassed by a model with a similar set of parameters. As with many phylogeographic methods, the accuracy of PHRAPL improves as the size of the dataset increases. While we report results for up to 100 loci, PHRAPL scales efficiently with genomic datasets because the gene trees estimated from 100s to 10,000s of loci are compared to the same simulated gene tree distributions to calculate the approximate likelihood. Performance decreases slightly when using gene trees estimated from simulated sequence data rather than the original genealogies, particularly in the case of isolation only datasets (Figs. S4-S5). Parameter estimates are generally accurate (Figs. S8-S9 and Table S3), with the exception of estimates of migration between ancestral populations, which were typically overestimated. We suspect that ancestral migration is difficult to estimate in general, as more recent events tend to erase this history.

**Fig. 2.**
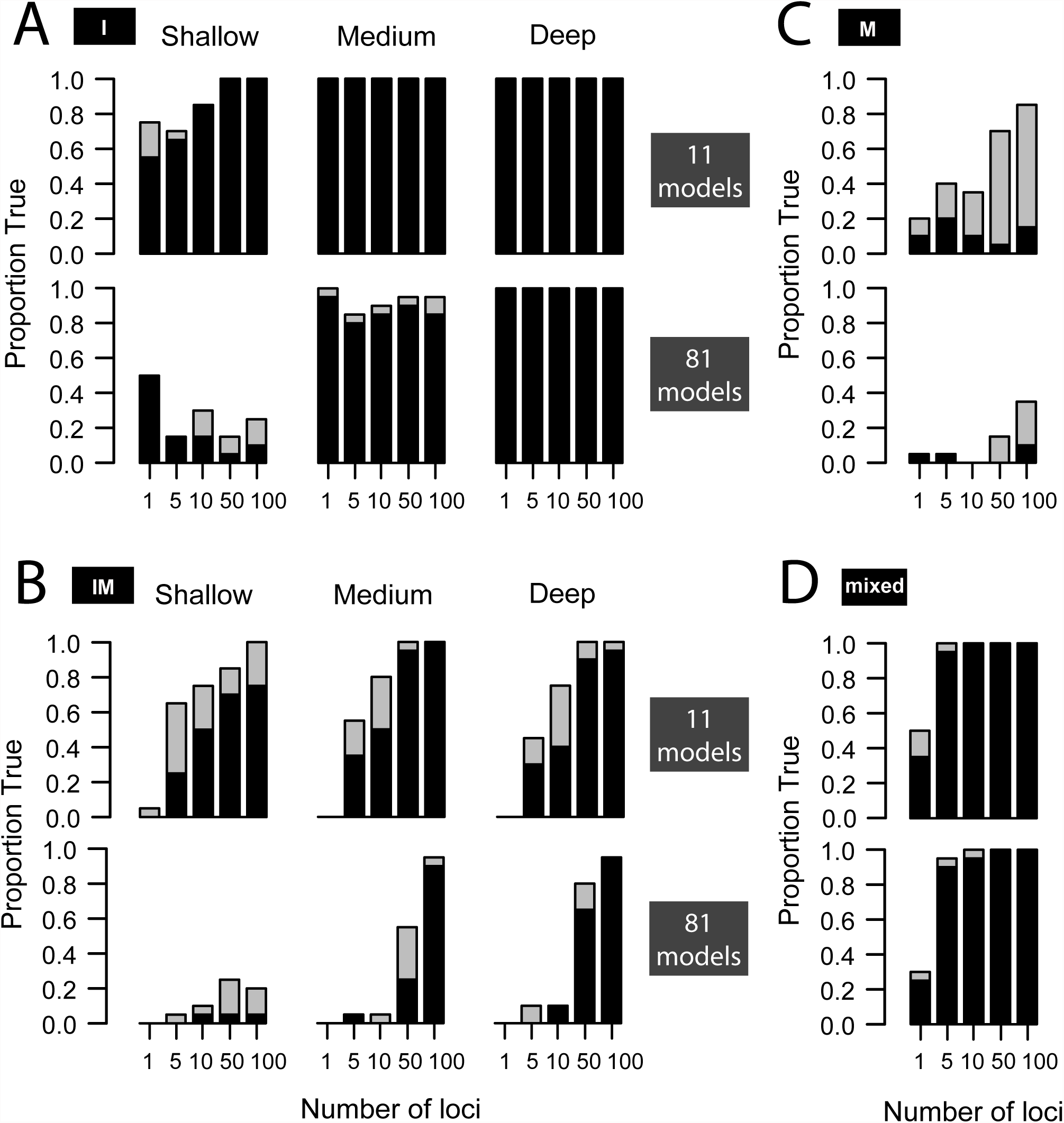
PHRAPL results from analysis of four types of simulated models (depicted in Fig. 1. A: isolation only; B isolation with migration; C: migration only; D: mixed). Results are shown for the entire model set (81 models) and for a reduced model set (11 models). Black bars give the proportion of 20 replicate analyses in which the true model garnered the highest AIC weight; grey bars give the proportion where the true model garnered the second highest AIC weight.

The number of models considered by a PHRAPL analysis also affects accuracy, particularly for IM models or IO models with shallow divergence, where accuracy declines with increases in the number of models considered by the analysis. Due to the nature of the model space, our large model set is predominately composed of IM models with subtle differences among migration matrices. Thus, with fewer loci or less time for lineage sorting, it becomes a challenge to consistently distinguish among models that include slight variations on the true migration scheme. We suggest that users evaluate parameter estimates in such cases. For example, although IM models were incorrectly inferred for many shallow isolation only datasets (although usually with the correct population topology; Fig. 2A), the estimated migration rates in these models were generally near zero (Fig. S8-S9 and Table S3). In practice, such a result would likely lead researchers (albeit in a circuitous way) to the correct inference (i.e., that migration is not very important for these datasets). The similarity in parameter values estimated using the small and large model sets (Fig. S8-S9) suggests that researchers do not necessarily need to identify the exact true model to accurately quantify the underlying processes. Rather, in keeping with the spirit of model-based inferences (e.g., Anderson 2008), it is perhaps best to view PHRAPL as a tool that is likely to identify the processes that have left a noticeable signal in the data. Parameters included in the models with the best AIC scores are those that reflect the dominant evolutionary processes that gave rise to the observed genetic patterns (Carstens et al. 2009).

The one scenario under which model inference was poor regardless of model set size was when the true model was an *n*-island model (Fig. 2C). In such a case, the models with the best AIC value usually included one or two divergence parameters in addition to full migration. This suggests that PHRAPL is biased towards IM models, which is worrisome for systems in which migration is so important as to have swamped out the genetic signal of the underlying divergence history. Phylogeography has been criticized for being overly reliant on tree thinking (Smouse 2008), and this aspect of PHRAPL’s performance should be improved. Nevertheless, high accuracy in migration rate estimates was still observed for these datasets (Fig. S8-S9), so it is unlikely that researchers would ignore migration in such systems altogether.

When parameters are estimated using model averaging, PHRAPL is precise in most cases (Fig. S8-S9). The exceptions are the timing of divergence and gene flow among ancestral lineages. In both cases, PHRAPL tends to overestimate these values, suggesting that genetic data contains less information about earlier processes than about more recent ones. In light of these results, PHRAPL is best suited to be used as a tool for identifying the optimal model for a given empirical system. Users should devote more effort to parameter estimation after an optimal model is identified, either by using a finer parameter grid within PHRAPL or by analyzing their data using an alternative method that implements the chosen model(s) in a full probabilistic framework.

Molecular systematists should be familiar with this relationship between model selection and parameter estimation; to estimate phylogenetic trees from sequence data researchers first use a program such as MODELTEST (Posada and Crandall 1998) to objectively choose a model of sequence evolution, and then other software to generate a precise estimate of the parameter in question (i.e., the phylogeny and branch lengths). However, it is worth noting that this workflow was not always in place. Prior to the widespread use of model selection, papers were routinely published that promoted unconventional relationships on the basis of phylogenetic trees that were likely poor estimates of the true parameter because they were estimated using inappropriate models (e.g., D’Erchia et al. 1996). In molecular systematics, parameters such as gamma for rate heterogeneity (Yang 1996) are important because they allow phylogeny (the true parameter of interest) to be estimated accurately, but are essentially nuisance parameters in terms of the inferences that result from the phylogeny estimate. In phylogeographic research, some parameters inherent to the models used to analyze the data are decidedly not nuisance parameters, particularly those that model evolutionary processes such as gene flow, genetic drift, or population history.

One criticism that could be made against PHRAPL is that it does not analyze all of the data. For example, it does not use raw gene sequences, nor does it use gene tree branch lengths. This is a compromise necessitated by its use of a discrete parameter to evaluate matches. In addition, it is often difficult to estimate a topology and branch lengths accurately using intraspecific data (Harding 1996), and thus branch length information would likely be noisy. Finally, there are the practical results: using just topologies, PHRAPL performs reasonably well in many cases. While we hope for methods in the future that can use more of the data, the results presented here suggest that topologies alone contain sufficient information to make inferences about important population-level evolutionary processes.

### Computation time required by PHRAPL

The computational time requirements of PHRAPL are similar to those of other methods commonly used by phylogeographers. Using a single core, individual models required 2.3 hours on average for the analysis of empirical data, resulting in a median time of 198 hours (8.2 computer-days) for the total model set (78 models). However, variance around this value was high (2.6 to 31 computer-days). Notably, PHRAPL is a method that is easy to implement in parallel because only the input data are shared between models, and substantial time savings can be accomplished by analyzing models across multiple cores on a single computer or split across several computers.

### Empirical studies

The reanalysis of recently published data suggests that the process of gene flow has been underappreciated by phylogeographic investigations. It is almost always implicated as an important evolutionary process by PHRAPL (Fig. 3 and Fig. S10), despite the fact that many of the original studies only considered species tree models (Table S2). Note that while the number of possible IM histories is inherently much larger than the number of MO and IO histories (90% of the model set was composed of IM models, whereas IO and MO models made up only 5% each), we normalized the model class probabilities depicted in Fig. 3 based on the frequency of each model class to account for this bias. The absence of support for isolation only models among empirical datasets suggests that reliance on species tree approaches (in our sample, the most commonly applied model to phylogeographic systems) may fail to account for important evolutionary processes. Moreover, parameter estimates of non-migration parameters can be affected when migration is ignored. For each of our 20 empirical datasets (including *Myotis*), there were four tree structures (polytomy and all three resolved trees) that were analyzed with and without migration. In 79 out of these 80 examples, the no-migration model had divergence times lower, often much lower, than the divergence times in the best fitting corresponding migration model. Over all the 80 pairs, the median branch length under a no-migration model was just 13% of the corresponding branch length for the best migration model. Thus, for empirical datasets, ignoring migration can have a significant effect on the resulting inference, even if it is not a parameter of interest *per se*.

**Fig. 3.**
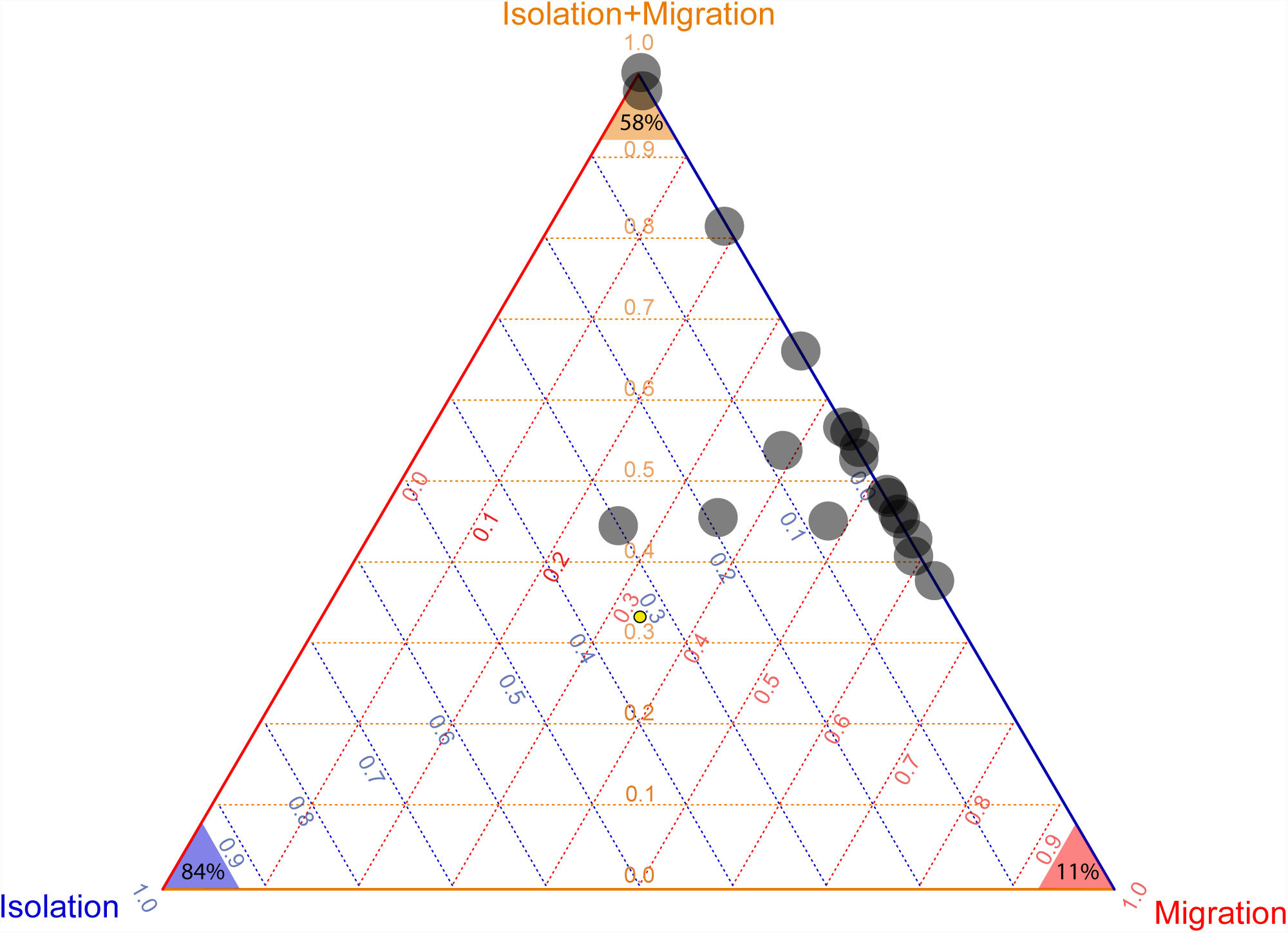
Triangle plot showing the weighted probabilities from PHRAPL analysis of 19 datasets. Each of the three vertices represents a commonly used approach to phylogeographic data analysis. The top corresponds to IMa2 (Hey 2010), the lower right to Migrate-n (Beerli and Felsenstein 2001) and the lower left to a species tree analysis (i.e., *BEAST; Heled and Drummond 2010). The probabilities of each analysis are shown decreasing from the respective vertex in increments of 0.1. Weighted probabilities were corrected for the unevenness of the model space in respect to the three model classes such that a dataset with equivalent probability for each of the models would appear in the center of the triangle (marked with a small yellow circle). Results indicate that there is very little support for the isolation-only model in these phylogeographic investigations. Percentages at the triangle tips give the proportion of the empirical studies that applied the corresponding model to the data.

Results from *Myotis lucifigus* mirror those seen in the analysis of the other empirical data. Of the 216 models included in the analysis, roughly 98.5 of the total model probability was represented by isolation-with-migration models (Table 1). The inferred topology matches the topology estimated using *BEAST (not shown), but gene flow is clearly a process that should be considered to understand the evolutionary history of this group. These results imply either that species delimitation approaches such as BPP (Yang and Ranalla 2010) and spedeSTEM (Ence and Carstens 2011) can accurately delimit lineages in the presence of moderate gene flow, or that they falsely delimit lineages by treating all shared polymorphism as the product of ancestral lineage sorting. Differentiating these scenarios would require a more nuanced understanding of both the timing of diversification and gene flow than can likely be provided by the PHRAPL analysis of the data analyzed here.

**Table 1.** Model selection results in *Myotis lucifugus*. Data from 4 subspecies (***<u>a</u>****lacensis*, ***<u>c</u>****arissima*, ***<u>l</u>****ucifugus*, ***<u>r</u>****elictus*) were analyzed using PHRAPL and 216 models. Shown are the model (letters with ’M’ represent models that included migration among the identified subspecies), the topology of the population tree, the number of parameters (*K*), the Akaike Information Criterion (Akaike 1973) score (AIC), and the model likelihood (Burnham and Anderson 1998). Subspecies are identified using the first letter of subspecies names.

| model | topology | K | AIC | wAIC |
| --- | --- | --- | --- | --- |
| c - l M | (((a,l)r)c) | 4 | 123.23 | 0.405 |
| c - r M | (((a,l)r)c) | 4 | 123.51 | 0.352 |
| a - c M | (((a,l)r)c) | 4 | 126.11 | 0.096 |
| a - c - l M | (((a,l)r)c) | 4 | 126.72 | 0.071 |
| a - l M | (((a,l)r)c) | 4 | 127.06 | 0.060 |
| species tree | (((a,l)r)c) | 3 | 129.87 | 0.015 |
| symmetric M | (((a,l)r)c) | 4 | 135.03 | 0.001 |

## Conclusion

Our reanalysis of 20 empirical datasets highlights the utility of phylogeographic model selection by demonstrating that the intuition of researchers (inclusive to some of the authors of this paper) is sometimes flawed in choosing the models used to analyze data from empirical systems. Optimal models for most datasets parameterize both gene flow and population divergence, suggesting that species tree methods (which do not consider gene flow) are over-simplifications for phylogeographic systems. Phylogeography has long been promoted as a ’bridge’ between population genetics and systematics (Avise et al. 1987), but the reliance on the species tree has constrained the field to one side of this continuum. For the first time, PHRAPL enables researchers to select demographic models without relying on their intuition about the processes likely to be important to their systems. By allowing a direct probabilistic assessment of nearly any coalescent model to the empirical data, PHRAPL represents a substantial addition to the methodological toolbox available to phylogeographers.

## Acknowledgements.

PHRAPL is funded by the National Science Foundation (DEB 1257669 / DEB-1257784). We thank the authors who generously provided empirical data. We thank Jack Sullivan, Darin Rokyta, members of the Carstens and O’Meara labs, as well as students in the first PHRAPL workshop for conversations related to this work.

## Data Archiving

Dryad once possible.

